# SynEM: Automated synapse detection for connectomics

**DOI:** 10.1101/099994

**Authors:** Benedikt Staffler, Manuel Berning, Kevin M. Boergens, Anjali Gour, Patrick van der Smagt, Moritz Helmstaedter

**Affiliations:** Department of Connectomics, Max Planck Institute for Brain Research, D-60438 Frankfurt, Germany; Biomimetic Robotics and Machine Learning, TUM, D-80333 Munich, Germany; current affiliation:Data Lab, VW Group, 80805 Munich, Germany

## Abstract

Nerve tissue contains a high density of chemical synapses, about 1 per µm^3^ in the mammalian cerebral cortex. Thus, even for small blocks of nerve tissue, dense connectomic mapping requires the identification of millions to billions of synapses, of which about 80-90% are excitatory synapses. While the focus of connectomic data analysis has been on neurite reconstruction, synapse detection becomes limiting when datasets grow in size and dense mapping is required. Here, we report SynEM, a method for automated detection of excitatory synapses from conventionally en-bloc stained 3D electron microscopy image stacks. The approach is based on a segmentation of the image data and focuses on classifying borders between neuronal processes as synaptic or non-synaptic. SynEM yields 98% precision and recall in binary excitatory cortical connectomes with no user interaction. It scales to large volumes of cortical neuropil, plausibly even whole-brain datasets. SynEM removes the burden of manual synapse annotation for large densely mapped connectomes.

## INTRODUCTION

The ambition to map neuronal circuits in their entirety has spurred substantial methodological developments in large-scale 3-dimensional microscopy (Denk & Horstmann, 2004, Hayworth et al., 2006, Knott et al., 2008, Eberle et al., 2015), making the acquisition of datasets as large as 1 cubic millimeter of brain tissue or even entire brains of small animals at least plausible (Mikula et al., 2012, Mikula & Denk, 2015). Data analysis, however, is lagging far behind (Helmstaedter, 2013). One cubic millimeter of gray matter in the mouse cerebral cortex, spanning the entire depth of the gray matter and comprising several presumed cortical columns (Fig. 1a), for example, contains at least 4 kilometers of axons, about 1 kilometer of dendritic shafts, about 1 billion spines (contributing an additional 2-3 kilometers of spine neck path length) and about 1 billion synapses (Fig. 1b). Initially, neurite reconstruction was so slow, that synapse annotation comparably paled as a challenge (Fig. 1c): when comparing the contouring of neurites (proceeding at 200-400 work hours per millimeter neurite path length) with synapse annotation by manually searching the volumetric data for synaptic junctions (Fig. 1d, proceeding at about 0.1 hour per µm^3^), synapse annotation consumed at least 20-fold less annotation time than neurite reconstruction (Fig. 1c). An alternative strategy for manual synapse detection is to follow reconstructed axons (Fig. 1e) and annotate sites of vesicle accumulation and postsynaptic partners. This axon-focused synapse annotation reduces synapse annotation time by about 8-fold for dense reconstructions (proceeding at about 1 min per potential contact indicated by a vesicle accumulation, which occurs every about 4-10 µm along axons in mouse cortex, Fig. 1e). Since in the mammalian cerebral cortex, 80-90% of synapses are excitatory, the detection of inhibitory synapses is about an order of magnitude less time consuming.

**Figure 1.**
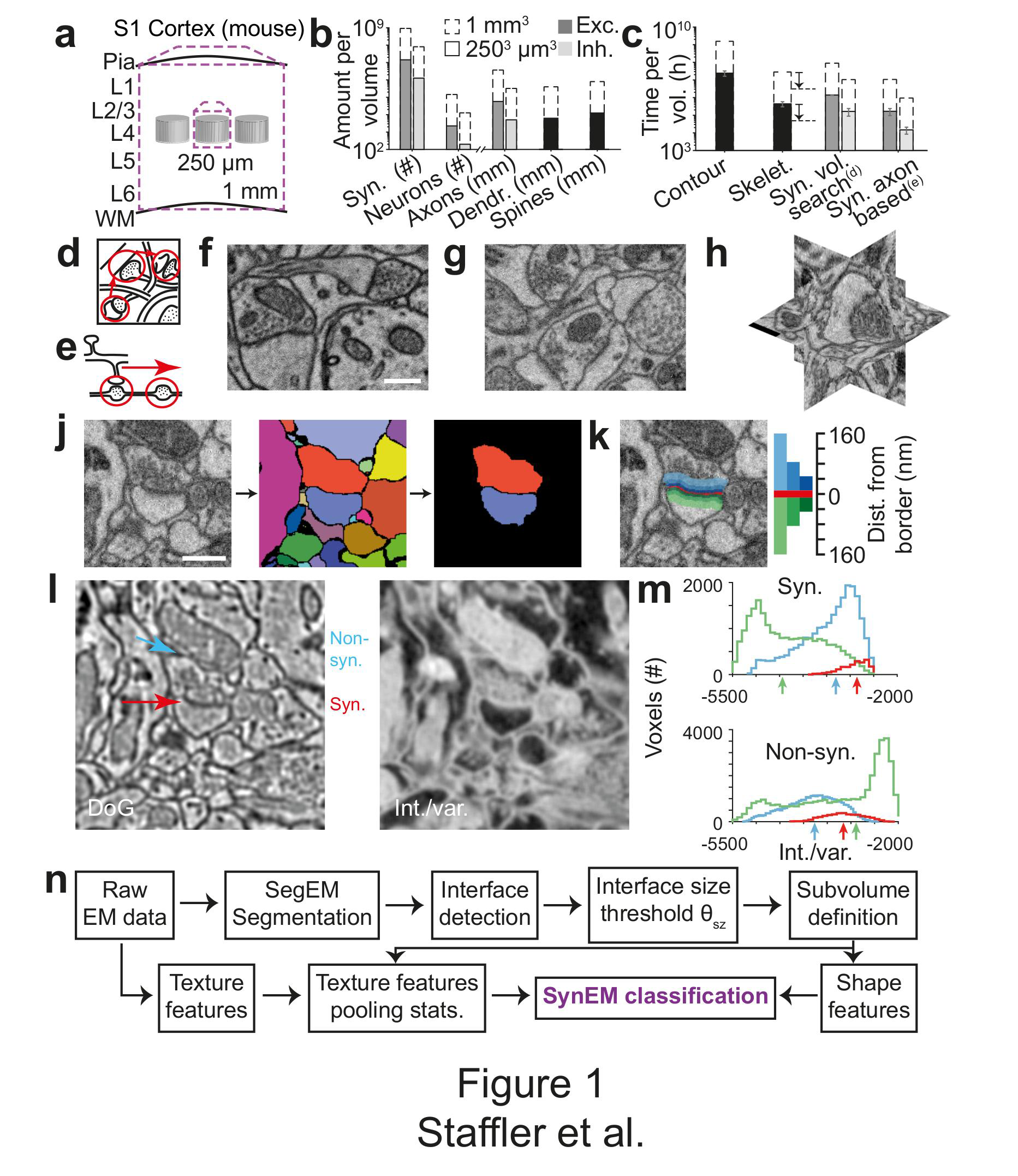
Synapse detection by classification of neurite interfaces. (**a**) Sketch of mouse primary somatosensory cortex (S1) with circuit modules (“barrels”) in cortical layer 4 and minimum required dataset extent for a “barrel” dataset (250 μm edge length) and a dataset extending over the whole cortical depth from pia to white matter (WM) (1 mm edge length). (**b**) Number of synapses and neurons, total axonal, dendritic and spine path length for the example datasets in (a) (White & Peters, 1993, Braitenberg & Schüz, 1998, Merchan-Perez et al., 2014). (**c**) Reconstruction time estimates for neurites and synapses; For synapse search strategies see sketches in d,e. Dashed arrows: latest skeletonization tools (webKnossos, Boergens et al., in prep.) allow for a further speed up of about 5-to-10-fold decreasing skeletonization effort below excitatory synapse detection time. (**d**) Volume search for synapses by visually investigating 3d image stacks and keeping track of already inspected locations takes about 0.1 h/μm3. (**e**) Axon-based synapse detection by following axonal processes and detecting synapses at boutons consumes about 1 min per bouton. (**f**) Examples of synapses imaged at an in-plane voxel size of 6 nm and (**g**) 12 nm in conventionally en-bloc stained and fixated tissue (Briggman et al., 2011, Hua et al., 2015). Note that synapse detection in high-resolution data is much facilitated in the plane of imaging. Large-volume image acquisition is operated at lower resolution, requiring better synapse detection algorithms. (**h**) Synapse with arbitrary orientation (shown in orthogonal planes). (**j**) Definition of interfaces used for synapse classification in SynEM. Raw EM data (left) is first volume segmented (using SegEM, Berning et al., 2015). Neighboring volume segments are identified (right). (**k**) Definition of perisynaptic subvolumes used for synapse classification in SynEM consisting of a border (red) and subvolumes adjacent to the neurite interface extending to distances of 40, 80 and 160 nm. (**l**) Example outputs of two texture filters: the difference of Gaussians (DoG) and the intensity/variance filter (int./var.). Note the clear signature of postsynaptic spine heads (right). (**m**) Distributions of int/var. texture filter output for image voxels at a synaptic (top) and non-synaptic interface (bottom). Medians over subvolumes are indicated (arrows, color scale as in k). (**n**) SynEM flow chart. Scale bars, 500 nm. Scale bar in f applies to g. Scale bar in j applies to j-l.

With the development of substantially faster annotation strategies for neurite reconstruction, however, the relative contribution of synapse annotation time to the total reconstruction time has substantially changed. Skeleton reconstruction (Helmstaedter et al., 2011) together with automated volume segmentations (Helmstaedter et al., 2013, Berning et al., 2015), allow to proceed at about 7-10 hours per mm path length (mouse retina, Helmstaedter et al., 2013) or 4-7 hours per mm (mouse cortex, Berning et al., 2015), thus about 50-fold faster than manual contouring. Recent improvements in online data delivery and visualization (webKnossos, Boergens, Berning et al., in revision) further reduce this by about 5-10 fold. Thus, synapse detection has become a limiting step in dense large-scale connectomics. Importantly, any further improvements in neurite reconstruction efficiency would be bounded by the time it takes to annotate synapses. Therefore, automated synapse detection for large-scale 3D EM data is critical.

High-resolution EM micrographs are the gold standard for synapse detection (Gray, 1959, Colonnier, 1968). Images acquired at about 2-4 nm in-plane resolution have been used to confirm chemical synapses using the characteristic intense heavy metal staining at the postsynaptic membrane, thought to be caused by the accumulated postsynaptic proteins (“postsynaptic density”, PSD), and an agglomeration of synaptic vesicles at the membrane of the presynaptic terminal. While synapses can be unequivocally identified in 2-dimensional images when cut perpendicularly to the synaptic cleft (Fig. 1f), synapses at oblique orientations or with a synaptic cleft in-plane to the EM imaging are hard or impossible to identify. Therefore, the usage of 3D EM imaging with a high resolution of 4-8 nm also in the cutting dimension (FIB/SEM, Knott et al., 2008) is ideal for synapse detection. For such data, automated synapse detection is available and successful (Kreshuk et al., 2011, Becker et al., 2012, 2013). However, FIB-SEM currently does not scale to large volumes required for connectomics of the mammalian cerebral cortex. Serial Blockface EM (Denk & Horstmann, 2004) scales to such mm^3^-sized volumes. However, SBEM provides a resolution just sufficient to follow all axons in dense neuropil and to identify synapses across multiple sequential images, independent of synapse orientation (the resolution typically is about 10x10x30 nm^3^; Fig. 1g). In this setting, synapse detection methods developed for high-in plane resolution data do not provide the accuracy required for fully automated synapse detection (see below). Here we report SynEM, an automated excitatory synapse detection method based on an automated segmentation of large-scale 3D EM data (using SegEM, Berning et al., 2015) from mouse cortex obtained using SBEM. SynEM is aimed at providing fully automated excitatory connectomes from large-scale EM data in which manual annotation or proof reading of synapses is not feasible. SynEM achieves precision and recall for single-synapse detection of 94% and 89%, and for binary neuron-to-neuron connectomes of 98% and 98% without any human interaction, essentially removing the synapse annotation challenge for large-scale excitatory mammalian connectomes.

## RESULTS

### Interface classification

We consider synapse detection as a classification of interfaces between neuronal processes as synaptic or non-synaptic (Fig. 1j; see also Mishchenko et al., 2010, Kreshuk et al., 2015, Huang et al., 2016). This approach relies on a volume segmentation of the neuropil sufficient to provide locally continuous neurite pieces (such as provided by SegEM, Berning et al., 2015, for SBEM data of mammalian cortex), for which the contact interfaces can be evaluated.

The unique features of excitatory synapses are distributed asymmetrically around the synaptic interface: presynaptically, large vesicle pools extend into the presynaptic terminal over at least 100-200 nm; postsynaptically, the PSD has a width of about 20-30 nm. To account for this surround information our classifier considers the subvolumes adjacent to the neurite interface explicitly and separately, unlike previous approaches (Kreshuk et al., 2015, Huang et al., 2016), up to distances of 40, 80, and 160 nm from the interface, restricted to the two segments in question (Fig. 1k; the interface itself was considered as an additional subvolume). We then compute a set of 11 texture features (Table 1, this includes the raw data as one feature), and derive 9 simple aggregate statistics over the texture features within the 7 subvolumes. In addition to previously used texture features (Kreshuk et al., 2011, Table 1), we use the local standard deviation, an intensity-variance filter and local entropy to account for the low-variance (“empty”) postsynaptic spine volume and presynaptic vesicle clouds, respectively (see Fig. 1l for filter output examples and Fig. 1m for filter distributions at an example synaptic and non-synaptic interface). The “sphere average” feature was intended to provide information about mitochondria, which often impose as false positive synaptic interfaces when adjacent to a plasma membrane. Furthermore, we employ 5 shape features calculated for the border subvolume and the two subvolumes extending 160 nm into the pre-and postsynaptic processes, respectively. Together, the feature vector for classification had 3224 entries for each interface (Table 1).

**Table: 1.** Overview of the classifier features used in SynEM, and comparison with existing methods. 11 3-dimensional texture filters employed at various filter parameters given in units of standard deviation (s) of Gaussian filters (s was 12/11.24 voxels in x and ydimension and 12/28 voxels in z-dimension, sizes of filters were set to σ/s*ceil(2*s)). When structuring elements were used, 1axbxc refers to a matrix of size a x b x c filled with ones and r specifies the semi-principal axes of an ellipsoid of length (r, r, r/2) voxels in x, y and zdimension. All texture features are pooled by 9 summary statistics (quantiles (0.25, 0.5, 0.75, 0, 1), mean, variance, skewness, kurtosis, respectively) over the 7 subvolumes around the neurite interface (see Fig. 1k). Shape features were calculated for three of the subvolumes: border (Bo) and the 160 nm distant pre-and postsynaptic volumes (160). Init. Class: initial SynEM classifier (see Fig. 2d for performance evaluation). N of instances: number of feature instances per subvolume (n = 7) and aggregate statistic (n = 9). *: Total number of employed features is 63 times reported instances for texture features. For shape features, the reported number is the total number of instances used, together yielding 3224 features total.

| Features | Kreshuk<br>et al.,<br>2011 | Becker<br>et al.,<br>2012 | Init.<br>Class. | Syn<br>EM | Parameters | N of<br>instances<br>* |
| --- | --- | --- | --- | --- | --- | --- |
| **Texture:** |  |  |  |  |  |  |
| Raw data |  | ✓ | ✓ | ✓ | - | 1 |
| 3 EVs of Structure<br>Tensor | ✓ | ✓ | ✓ | ✓ | $(\sigma_w, \sigma_d) = \{(s,s), (s,2s), (2s,s), (2s,2s), (3s,3s)\}$ | 15 |
| 3 EVs of Hessian | ✓ | ✓ | ✓ | ✓ | $\sigma = \{s, 2s, 3s, 4s\}$ | 12 |
| Gaussian Smoothing | ✓ | | ✓ | ✓ | $\sigma = \{s, 2s, 3s\}$ | 3 |
| Difference of Gaussians | ✓ | | | ✓ | $(\sigma,k) = \{(s, 1.5), (s, 2), (2s, 1.5), (2s, 2), (3s, 1.5)\}$ | 5 |
| Laplacian of Gaussian | ✓ | ✓ | ✓ | ✓ | $\sigma = \{s, 2s, 3s, 4s\}$ | 4 |
| Gauss Gradient Magn. | ✓ | ✓ | ✓ | ✓ | $\sigma = \{s, 2s, 3s, 4s, 5s\}$ | 5 |
| Local standard deviation | | | | ✓ | $U = 1_{5 \times 5 \times 5}$ | 1 |
| Int./var. | | | | ✓ | $U = \{1_{3 \times 3 \times 3}, 1_{5 \times 5 \times 5}\}$ | 2 |
| Local entropy | | | | ✓ | $U = 1_{5 \times 5 \times 5}$ | 1 |
| Sphere average | | | | ✓ | $r = \{3, 6\}$ | 2 |
| **Shape:** |  |  |  |  |  |  |
| Number of voxels |  |  | ✓ | ✓ | Bo, 160 | 3 |
| Diameter (vx based) |  |  |  | ✓ | Bo | 1 |
| Lengths of principal axes |  |  |  | ✓ | Bo | 3 |
| Principal axis product |  |  |  | ✓ | 160 | 1 |
| Convex hull (vx based) |  |  |  | ✓ | Bo, 160 | 3 |

### SynEM workflow and training data

We developed and tested SynEM on a dataset from layer 4 (L4) of mouse primary somatosensory cortex (S1) acquired using SBEM (dataset ex145_07x2, Boergens et al., in prep.; the dataset was also used in developing SegEM, Berning et al., 2015). The dataset had a size of 93 x 60 x 93 µm^3^ imaged at a voxel size of 11.24 x 11.24 x 28 nm^3^. The dataset was first volume segmented (SegEM, Berning et al., 2015, Fig. 1j, see Fig. 1n for a SynEM workflow diagram). Then, all interfaces between all pairs of volume segments were determined, and the respective subvolumes were defined. Next, the texture features were computed on the entire dataset and aggregated as described above. Finally, the shape features were computed. Then, the SynEM classifier was implemented to output a synapse score for each interface and each of the two possible pre-to-postsynaptic directions (Fig. 2a-c). The SynEM score was then thresholded to obtain an automated binary classification of interfaces into synaptic / non-synaptic (θ in Fig. 2a). Since the SynEM scores for the two possible synaptic directions were rather disjunct in the range of relevant thresholds (Fig. 2b), we used the larger of the two scores for classification. By introducing a second threshold θ_2_ on the SynEM score, one can use SynEM instead in a semi-automated setting where a range of interfaces (between θ_1_ and θ_2_ Suppl. Fig. 1b) is manually inspected to further improve precision and recall. In the following, we report results on the fully-automated version of SynEM (Fig. 2d, e).

**Figure 2.**
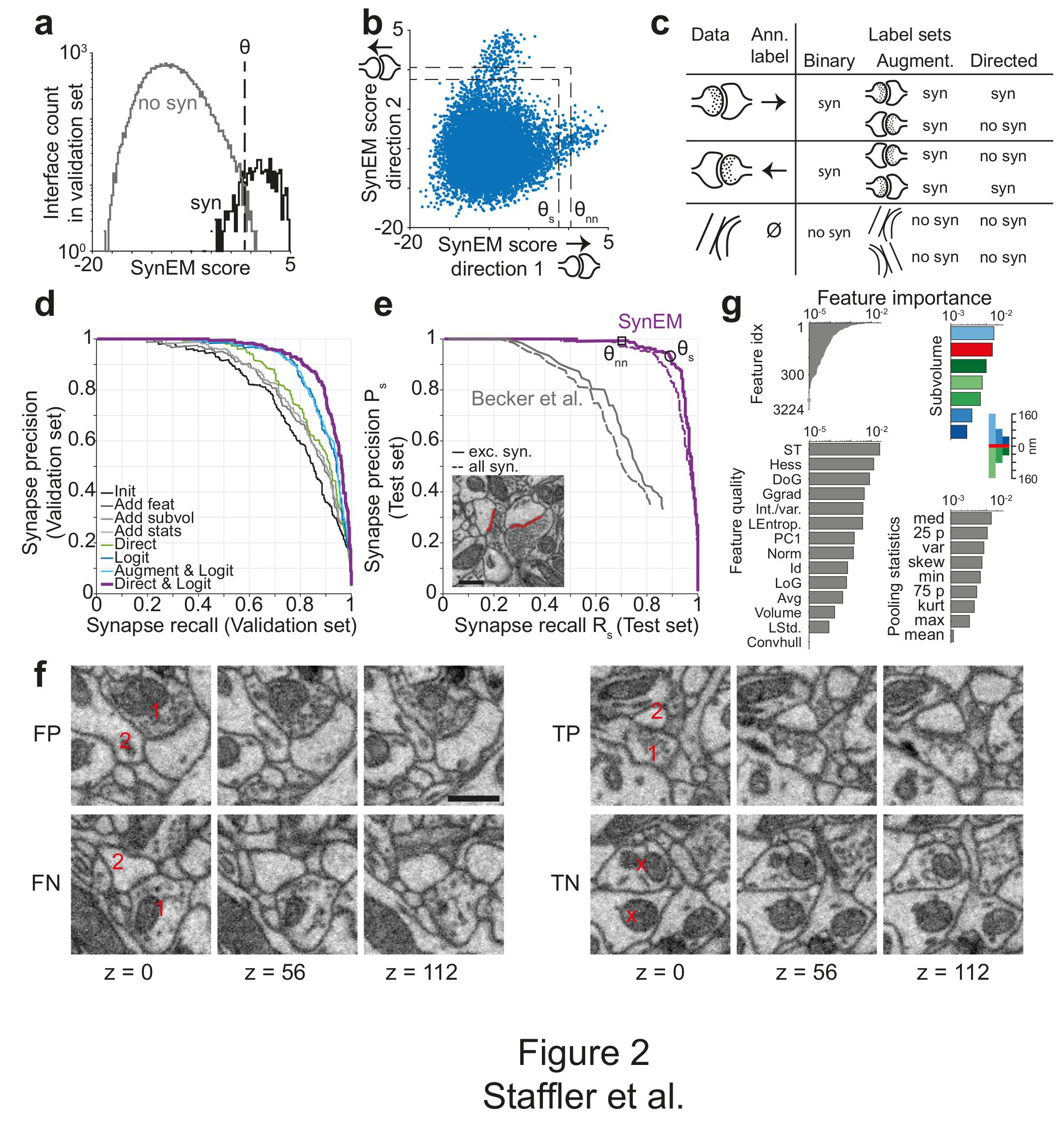
SynEM training and evaluation. (**a**) Histogram of SynEM scores calculated on the validation set. Fully automated synapse detection by thresholding the SynEM score at threshold θ. (**b**) SynEM scores for the two possible directions of interfaces. Note that SynEM scores are disjunct in a threshold regime used for best single synapse performance (θs) and best neuron-to-neuron recall and precision (θnn), see Fig. 3, indicating a clear bias towards one of the two possible synaptic directions. (**c**) Strategy for label generation. Based on annotator labels (Ann. Label), three types of label sets were generated: Initial label set ignored interface orientation (Binary); Augmented label set included mirror-reflected interfaces (Augment.); Directed label set used augmented data but considered only one synaptic direction as synaptic (Directed). (**d**) Classification performance for different features, aggregation statistics, classifier parameters and label sets. Init: initial classifier used (see Table 1). The initial classifier was extended by using additional features (Add feat, see Table 1, first row), 40 and 80 nm subvolumes for feature aggregation (Add subvol, see Fig. 1k) and aggregate statistics (Add stats, see Table 1). Logit: Classifier trained on full feature space using LogitBoost. Augment & Logit: Logit classifier trained on augmented label set (see Fig. 2c). Direct & Logit: Logit classifier trained on directed label set (see Fig. 2c). (**e**) Test set performance of SynEM (purple) and the synapse detection algorithm of (Becker et al., 2012, gray) for spine and shaft synapses and for spine synapses, only. Inset shows example test set labels. Threshold values for optimal single synapse detection performance (black circle) and an optimal connectome reconstruction performance (black square, see Fig. 3). (**f**) Correct and incorrect classification examples obtained from SynEM classification at θs (circle in e), 3 image planes spaced by 56 nm are shown (i.e. every second SBEM data slice). Note that detection in single or a few images is challenging in data of this resolution. Only fully 3-dimensional treatment allows high-accuracy detection. 1: presynaptic; 2: postsynaptic; x: non-synaptic (**g**) Ranked importance of SynEM features for classification. All features (top left), relevance of feature quality (bottom left), subvolumes (top right) and pooling statistics (bottom right). Note that only 378 features contribute to classification. See Table 3 for the 10 highest importance feature instances, Table 1 for feature name abbreviations, and text for details. Scale bars, 500 nm.

We obtained labels for SynEM training and validation by presenting raw data volumes of (1.6 x 1.6 x 0.7-1.7) µm^3^ that surrounded the segment interfaces to trained student annotators (using a custom-made annotation interface in Matlab, Suppl. Fig. 1a). The raw data was rotated such that the interface was most vertically oriented in the image plane presented to the annotators; the two interfacing neurite segments were colored transparently for identification (this could be switched off by the annotators when inspecting the synapse, see Methods for details). Annotators were asked to categorize the presented interface as either non-synaptic, pre-to-postsynaptic, or post-to-presynaptic (Fig. 2a, Suppl. Fig. 1a). The synaptic labels were then verified by an expert neuroscientist. A total of 75,383 interfaces (1,858 synaptic, 73,525 non-synaptic) were annotated in image volumes drawn from 40 locations within the entire EM dataset. About 80% of the labels (1467 synaptic, 61,619 non-synaptic) were used for training, the remaining were used for validation. Initially, we interpreted the annotator’s labels in a binary fashion: irrespective of synapse direction, the label was interpreted as synaptic (and non-synaptic otherwise, Fig. 2c, “binary”). We then augmented the training data by including mirror-reflected copies of the originally presented synapses, maintaining the labels as synaptic (irrespective of synapse direction) and non-synaptic (Fig. 2c, “augmented”). Finally, we changed the labels of the augmented training data to reflect the direction of synaptic contact: only synapses in one direction were labeled as synaptic, and non-synaptic in the inverse direction (Fig. 2c “directed”).

Fig. 2d shows the effect of the choice of features, aggregate statistics, classifier parameters and label types on SynEM precision and recall. Our initial classifier used the texture features from Kreshuk et al., 2011 with minor modifications and in addition the number of voxels of the interface and the two interfacing neurite segmentation objects (restricted to 160 nm distance from the interface) as a first shape feature (Table 1). This classifier provided only about 70% precision and recall (Fig. 2d). We then extended the feature space by adding more texture features capturing local image statistics (Table 1) and shape features. In particular, we added filters capturing local image variance in an attempt to represent the “empty” appearance of postsynaptic spines, and the presynaptic vesicle clouds imposing as high-frequency high-variance features in the EM images. Also, we added more subvolumes over which features were aggregated (see Fig. 1k), increasing the dimension of the feature space from 603 to 3224. Together with additional aggregate statistics, the classifier reached about 75% precision and recall. A substantial improvement was obtained by switching from an ensemble of decision-stumps (one-level decision tree) trained by AdaBoostM1 (Freund & Schapire, 1997) as classifier to decision stumps trained by LogitBoost (Friedman et al., 2000). In addition, the directed label set proved to be superior. Together, these improvements yielded a precision and recall of 87% and 86% on the validation set (Fig. 2d).

We then evaluated the best classifier from the validation set (Fig. 2d, ‘Direct & Logit’) on a separate test set. This test set was a dense volume annotation of all synapses in a randomly positioned region containing dense neuropil of size 5.8 x 5.8 x 7.2 µm^3^ from the L4 mouse cortex dataset. All synapses were identified by 2 experts, which included the reconstruction of all local axons, and validated once more by another expert on a subset of synapses. In addition we marked all shaft (to a large majority inhibitory) synapses in the test set and evaluated our approach for all 235 synapses in the test set and for 204 spine (to a large majority excitatory) synapses only (Fig. 2e). We compared SynEM to the synapse detection algorithms from Kreshuk et al.,2011, Becker et al., 2012 and Kreshuk et al., 2014 on the test set (Fig. 2e, only best result from Becker et al., 2012 shown, see Methods for training details). Especially in the high recall regime SynEM was able to significantly outperform previous approaches on the SBEM dataset with its lower resolution (see also Suppl. Table 1 for a comparison of detection methods). SynEM achieves precision and recall rates for single excitatory synapse detection of 94% and 89%, respectively (at SynEM score threshold θ_s_, black circle in Fig. 2e).

Fig. 2f shows examples of correct and incorrect SynEM classification results (evaluated at θ_s_). Typical sources of errors are vesicle clouds close to membranes that target nearby neurites (Fig. 2f, FP), Mitochondria in the pre-and/or postsynaptic process, very small vesicle clouds and/or small PSDs (Fig. 2f, FN), and remaining SegEM segmentation errors (see Suppl. Material for 3-dimensional image sequences of TP, FP, TN, FN calssifications).

We then asked which features had highest classification power, and whether the newly introduced texture and shape features contributed to classification. Boosted decision-stump classifiers allow the ranking of features according to their classification importance (Fig. 2g). 378 out of 3224 features contributed to classification (leaving out the remaining features did not reduce accuracy, thus allowing us to reduce computation time). The 10 features with highest discriminative power (Table 2) in fact contained two of the added texture filters (int-var and local entropy) and a shape feature. The three most distinctive subvolumes (Fig. 2g) were the large presynaptic subvolume, the border and the small postsynaptic subvolume. This suggests that the asymmetry in pre- vs. postsynaptic aggregation volumes in fact contributed to classification performance, with a focus on the presynaptic vesicle cloud and the postsynaptic density.

**Table 2:** SynEM features ranked by ensemble predictor importance. See Fig.1 and Methods for details. Note that two of the newly introduced features and one of the shape features had high classification relevance (Local entropy, Int./var., Principal axes length; cf. Table 1).

| Rank | Feature | Parameters | Subvolume | Aggregate statistic |
| --- | --- | --- | --- | --- |
| 1 | EVs of Struct. Tensor (largest) | $\sigma_w = 2s$,<br>$\sigma_D = s$ | 160 nm, S1 | Median |
| 2 | EVs of Struct. Tensor (smallest) | $\sigma_w = 2s$,<br>$\sigma_D = s$ | 160 nm, S1 | Median |
| 3 | Local entropy | $U = 1_{5 \times 5 \times 5}$ | 160 nm, S2 | Variance |
| 4 | Difference of Gaussians | $\sigma = 3s$,<br>$k = 1.5$ | Border | 25<sup>th</sup> perc |
| 5 | Difference of Gaussians | $\sigma = 2s$,<br>$k = 1.5$ | Border | Median |
| 6 | EVs of Struct. Tensor (middle) | $\sigma_w = 2s$,<br>$\sigma_D = s$ | 40 nm, S2 | Min |
| 7 | Int./var. | $U = 1_{3 \times 3 \times 3}$ | Border | 75<sup>th</sup> perc |
| 8 | EVs of Struct. Tensor (largest) | $\sigma_w = 2s$,<br>$\sigma_D = s$ | 80 nm, S1 | 25<sup>th</sup> perc |
| 9 | Gauss gradient magnitude | $\sigma = s$ | 40 nm, S2 | 25<sup>th</sup> perc |
| 10 | Principal axes length (2nd) | - | Border | - |

### SynEM for connectomes

We so far evaluated SynEM on the basis of the detection performance of single synaptic interfaces. Since we are interested in measuring the connectivity matrices of mammalian cortical circuits (connectomes) we obtained a statistical estimate of connectome error rates based on synapse detection error rates. We assume that the goal is a binary connectome containing the information whether pairs of neurons are connected or not. Automated synapse detection provides us with weighted connectomes reporting the number of synapses between neurons, from which we can obtain binary connectomes by considering all neuron pairs with at least γ_nn_ synapses as connected (Fig. 3a). Synaptic connections between neurons in the mammalian cerebral cortex have been found to be established via multiple synapses per neuron pair (Fig. 3b, Feldmeyer et al., 1999, Feldmeyer et al., 2002, Feldmeyer et al., 2006, Frick et al., 2008, Markram et al., 1997, ranges 1-8 synapses per connection, mean 4.3 ± 1.4). The effect of synapse recall R_s_ on recall of neuron-to-neuron connectivity R_nn_ can be estimated (Fig. 3c) for each threshold γ_nn_ given the distribution of the number of synapses per connected neuron pair n_syn_. For connectomes in which neuron pairs with at least one detected synapse are considered as connected (γ_nn_= 1), a neuron-to-neuron connectivity recall R_nn_ of 98% can be achieved with a synapse detection recall R_s_ of 70.6% (Fig. 3c, black arrow) if synapse detection is independent between multiple synapses of the same neuron pair. SynEM achieves 99.3% synapse detection precision P_s_ at this recall (Fig. 2e).

**Figure 3.**
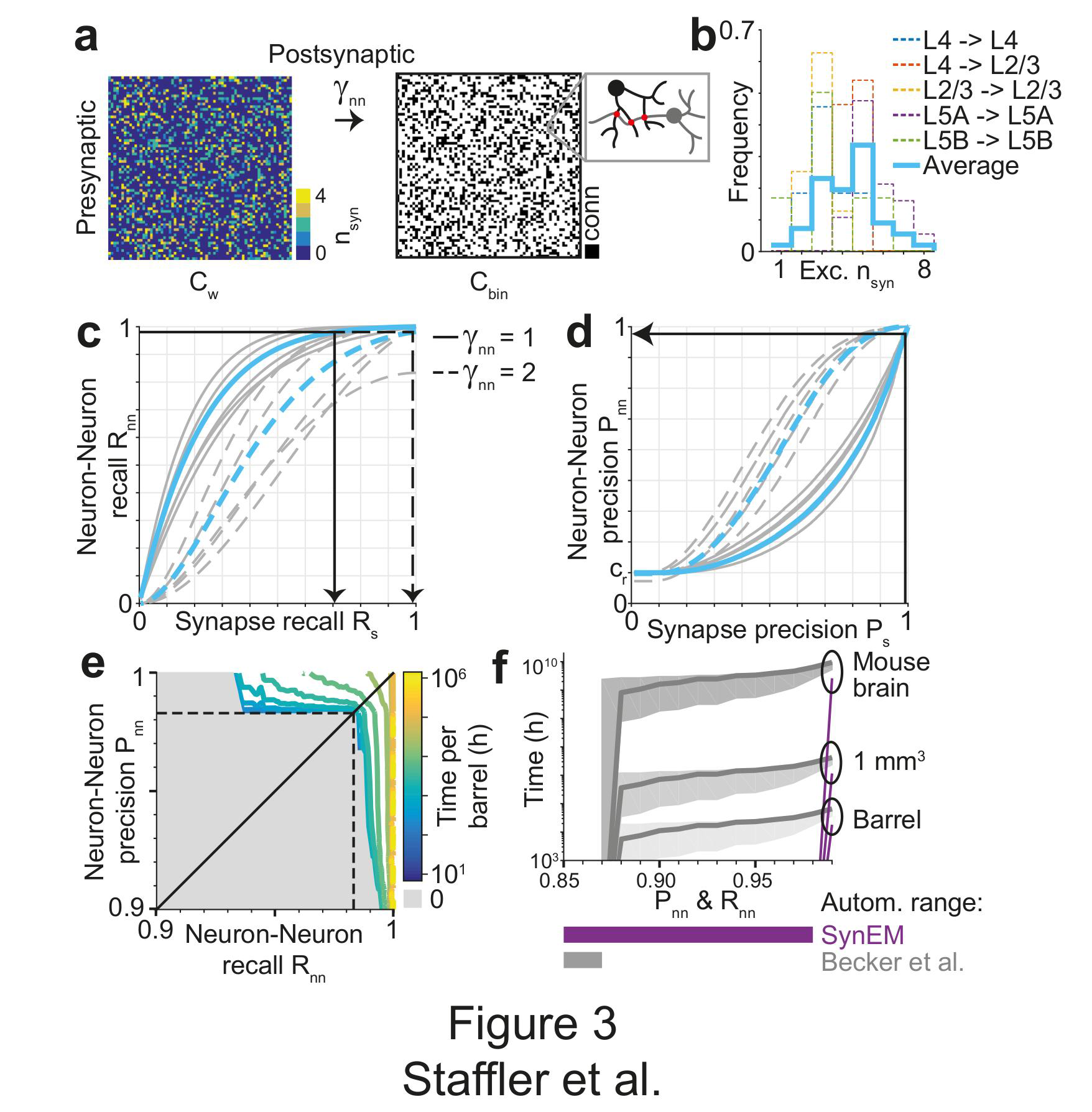
SynEM for connectomes. (**a**) Sketch of a weighted connectome (left) reporting the number of synapses per neuron-to-neuron connection, transformed into a binary connectome (middle) by considering neuron pairs with at least γnn synapses as connected. (**b**) Distribution of synapse number for connected excitatory neuron pairs obtained from paired recordings in rodent cerebral cortex (Feldmeyer et al., 1999, Feldmeyer et al., 2002, Feldmeyer et al., 2006, Frick et al., 2008, Markram et al., 1997). Average distribution (cyan) is used for the precision estimates in the following (see Suppl. Table 2). (**c**) Relationship between SynEM recall for single interfaces (synapses) Rs and the ensuing neuron-to-neuron connectome recall Rnn (recall in Cbin, a) for each of the excitatory cortico-cortical connections (summarized in b) and for connectome binarization thresholds of γnn = 1 and γnn = 2 (full and dashed, respectively). (**d**) Relationship between SynEM precision for single interfaces (synapses) Ps and the ensuing neuron-to-neuron connectome precision Pnn. Colors as in c. (**e**) Regime of neuron-to-neuron precision Pnn and recall Rnn that can be obtained fully automatically by SynEM (gray). Up to 98% precision Pnn and recall Rnn no manual interaction is required. (**f**) Comparison of fully automated classification regime for SynEM (purple) and the best competing method (Becker et al., 2012, gray). Note that SynEM shifts the fully automated regime to values which typically exceed neuronal wiring noise (below 2% error rates), allowing for fully automated synapse detection.

The resulting precision of neuron-to-neuron connectivity P_nn_ then follows from the total number of synapses in the connectome 
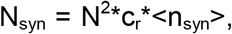
 with c_r_ the pairwise connectivity rate, about 20% for local connections in cortex (Feldmeyer et al., 1999), <n_syn_> the mean number of synapses per connection (4.3 ± 1.4, Fig. 3b), and N^2^ the size of the connectome. A fraction R_s_ of these synapses is detected (true positive detections, TPs). The number of false positive (FP) synapse detections was deduced from TP and the synapse precision P_s_ as FP=TP*(1-P_s_)/P_s_, yielding R_s_*N_syn_*(1-P_s_)/P_s_ false positive synapse detections. These we assumed to be distributed randomly on the connectome and estimated how often at least γ_nn_ synapses fell into a previously empty connectome entry. These we considered as false positive connectome entries, whose rate yields the binary connectome precision P_nn_ (see Methods for details of the calculation). At R_nn_ of 98%, SynEM yields a neuron-to-neuron connection precision P_nn_ of 98% (Fig. 3d, black arrow, Fig. 3e). Error rates of 2% for missed connections and for wrongly detected connections are well below the noise of synaptic connectivity so far found in real biological circuits (e.g., Helmstaedter et al., 2013, Bartol et al., 2015), and thus likely sufficient for a large range of studies involving the mapping of cortical connectomes.

In summary, SynEM provides fully automated synapse detection for binary mammalian connectomes up to 98% precision and recall, a range which was previously prohibitively expensive to attain by existing methods (Fig. 3f).

### Local cortical connectome

We applied SynEM to a local cortical connectome between 100 axons and 100 postsynaptic processes in the dataset from L4 of mouse cortex (Fig. 4a, b). We first detected all contacts and calculated the total contact area between each pair of pre-and postsynaptic processes (“contactome”, Fig. 4c). We then classified all contacts using SynEM (at the classification threshold θ_nn_ (Table 3) yielding 98% neuron-to-neuron precision and recall) followed by binarization at γ_nn_ = 1 to obtain the binary connectome C_bin_ (Fig. 4d).

**Table 3:** SynEM score thresholds and associated precision and recall. SynEM score chosen for best single synapse performance (θs) and a neuron-to-neuron connection recall and precision of 98% (θnn) with corresponding single synapse precision and recall rates (Ps, Rs, respectively) and estimated neuron-to-neuron precision and recall rates (Pnn, Rnn, respectively) for connectome binarization thresholds of nn = 1 and nn = 2.

| Threshold score | Single synapse $P_s/R_s$ | Neuron-to-neuron $P_{nn}/R_{nn}$ | |
| --- | --- | --- | --- |
| | | $\gamma_{nn} = 1$ | $\gamma_{nn} = 2$ |
| $\theta_s = -1.23$ | 94.3% / 88.7% | 84.6% / 99.7% | 99.6% / 95.8% |
| $\theta_{nn} = 0.03$ | 99.3% / 70.6% | 98.3% / 98.1% | 100% / 87.4% |

**Figure 4.**
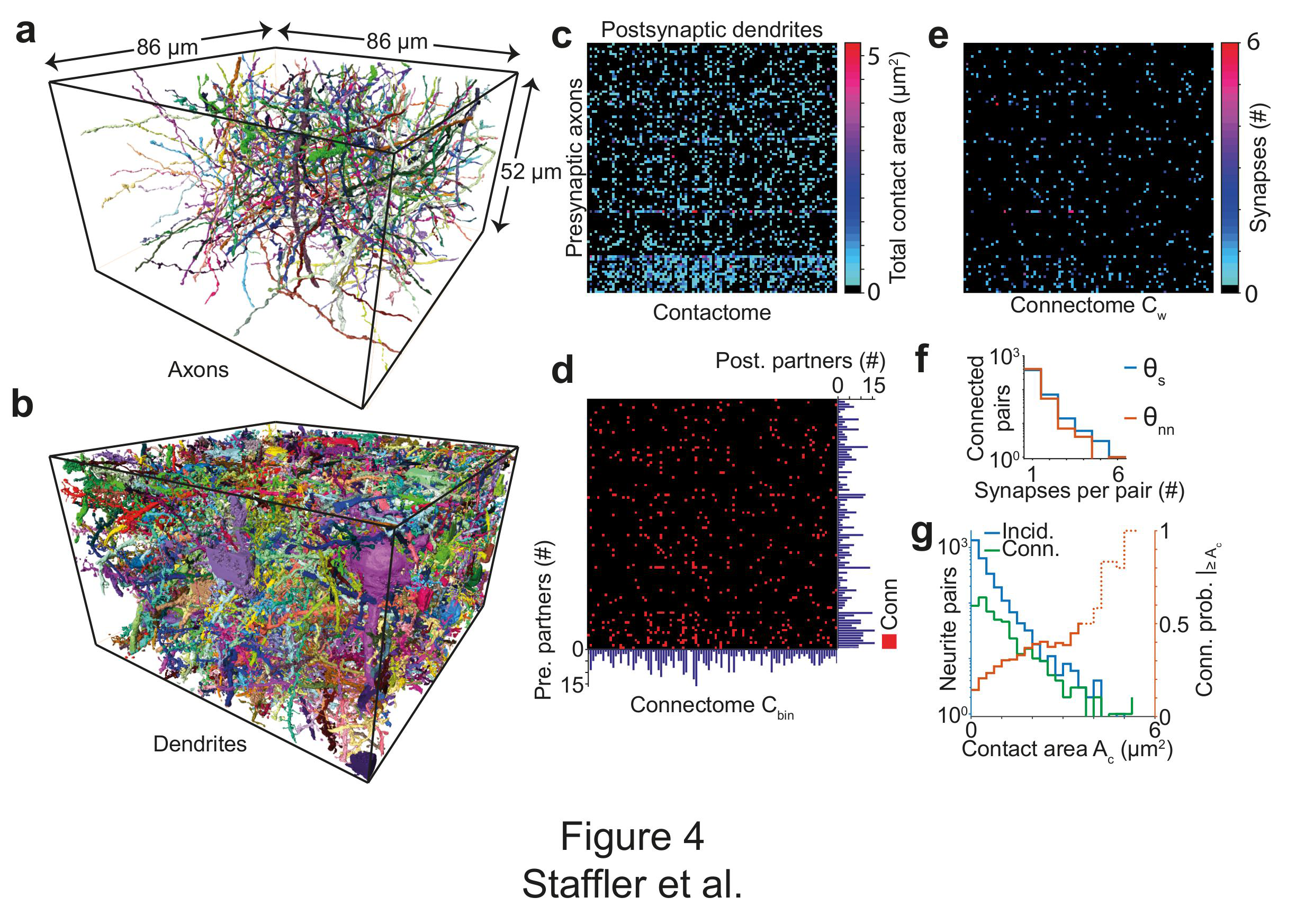
SynEM for mapping a local cortical connectome. (**a, b**) 100 axonal and dendritic processes within a volume sized 86 x 52 x 86 μm3 from layer 4 of mouse cortex. (**c**) Contactome reporting total contact area between pre-and postsynaptic processes. (**d**) Binary connectome obtained at the SynEM threshold θnn yielding neuron-to-neuron recall and precision of 98% (see Fig 2e, black square). Histograms of the number of pre (right)-and postsynaptic (bottom) partners for each process. (**e**) Weighted connectome reporting the number of synapses obtained at the SynEM threshold optimal for single synapse detection θs (Fig 2e, black circle); SynEM was evaluated only for neuron pairs detected as connected in the binary connectome (d). (**f**) Histogram of the number of synapses between connected neurites shown for the single-synapse optimized (θs, blue) SynEM threshold. For comparison, the result from the connectome-optimized SynEM threshold is also shown (θnn, red). (**g**) Total contact area for connected (green) and incidentally touching (blue) neurite pairs. Fraction of neurite pairs that were synaptically coupled and exceeded a certain total contact area Ac (red; full line for up to 99.5% of connections with non-zero contact area, dashed for remaining 0.5%). Note that this fraction does not exceed 50% over a wide range of contact areas, unlike in mouse retina (Fig. 3e in Helmstaedter et al., 2013).

In order to obtain an improved weighted connectome C_w_ (Fig. 4e), we finally applied SynEM to those contacts belonging to neurite pairs indicated as connected in the binary connectome. Here we used SynEM at the classification threshold θ_s_ yielding 89% single synapse recall and 94% single synapse precision. The detected synapses were clustered when they were closer than 1500 nm for a given neurite pair. This allowed us to concatenate large synapses with multiple active zones or multiple contributing SegEM segments into one (Suppl. Fig. 1d).

The resulting connectome contained 608 synapses distributed over 474 connections (Fig. 4e, f). We finally determined whether total contact area was predictive of synaptic connectivity for these connections in cortex (Fig. 4g). Note that while in the retina (Fig. 3e in Helmstaedter et al., 2013), total contact of more than about 1 µm^2^ was highly predictive of synaptic contact, in cortex such prediction is not possible (see also Mishchenko et al., 2010, Bock et al., 2011, Kasthuri et al., 2015, Lee et al., 2016).

## DISCUSSION

We report SynEM, a toolset for automated synapse detection in connectomics. The particular achievement is that the synapse detection for densely mapped excitatory connectomes from the mammalian cerebral cortex is fully automated yielding below 2% residual error in the connectome. With this, synapse detection is removed as a bottleneck in large-scale mammalian connectomics.

Evidently, synapse detection is facilitated in high-resolution EM data, and becomes most feasible in FIB-SEM data at a resolution of about 4-8 nm isotropic (Kreshuk et al., 2011). Yet, only by compromising resolution for speed (and thus volume) of imaging, the mapping of large, potentially even whole-brain connectomes is becoming plausible. Therefore it was essential to obtain automated synapse detection for EM data that is of lower resolution, but actually is scalable to such volumes.

In addition to high image resolution, synapse detection is substantially simplified for human annotators when employing recently proposed special fixation procedures that enhance the extracellular space in 3D EM data (Pallotto et al., 2015). In such data, direct touch between neurites has a very high predictive power for the existence of a (chemical or electrical) synapse, since otherwise neurite boundaries are separated by extracellular space. Thus, it is expected that such data substantially simplifies automated synapse detection. The advantage of SynEM is that it achieves fully automated excitatory synapse detection in conventionally stained and fixated 3D EM data, in which neurite contact is frequent. Such data is widely used, and acquiring such data does not require special fixation protocols.

Together, SynEM resolves excitatory synapse detection for high-throughput cortical connectomics of mammalian brains, making the efficiency of neurite reconstruction again the bottleneck for connectomic analysis.

## METHODS

### Annotation time estimates

Neuropil composition (Fig. 1b) was considered as follows: Neuron density of 157,500 per mm^3^ (White & Peters, 1993), axon path length density of 4 km per mm^3^ and dendrite path length density of 1 km per mm^3^ (Braitenberg & Schüz, 1998), spine density of about 1 per µm dendritic shaft length, with about 2 µm spine neck length per spine (thus twice the dendritic path length), synapse density of 1 synapse per µm^3^ (Merchan-Perez et al., 2014) and bouton density of 0.1 – 0.25 per µm axonal path length (Braitenberg & Schüz, 1998). Annotation times were estimated as 200 - 400 h per mm path length for contouring, 3.7 – 7.2 h/mm path length for skeletonization (Helmstaedter et al., 2011, Helmstaedter et al., 2013, Berning et al., 2015), 0.6 h/mm for flight-mode annotation (Boergens, Berning et al., in revision), 0.1 h/µm^3^ for synapse annotation by volume search (estimated form the test set annotation) and an effective interaction time of 60 s per identified bouton for axon-based synapse search. All annotation times refer to single-annotator work hours, redundancy may be increased to reduce error rates in neurite and synapse annotation in these estimates (see Helmstaedter et al., 2011).

### EM image dataset and segmentation

SynEM was developed and tested on a SBEM dataset from layer 4 of mouse primary somatosensory cortex (dataset 2012-09-28_ex145_07x2, K. M. B. and M. H., unpublished data, see also Berning et al., 2015). Tissue was conventionally en-bloc stained (Briggman et al., 2011) with standard chemical fixation yielding compressed extracellular space (compare to Pallotto et al., 2015).

The image dataset was volume segmented using the SegEM algorithm (Berning et al., 2015). Briefly, SegEM was run using CNN 20130516T204040_8,3_ and segmentation parameters as follows: r_se_ = 0; θ_ms_ = 50; θ_hm_ = 0.39; (see last column in Table 2 in (Berning et al., 2015)). For training data generation, a different voxel threshold for watershed marker size θ_ms_ = 10 was used. For test set and local connectome calculation the SegEM parameter set optimized for whole cell segmentations was used (r_se_ = 0; θ_ms_ = 50; θ_hm_ = 0.25, see Table 2, Berning et al., 2015).

### Neurite interface extraction and subvolume definition

Interfaces between a given pair of segments in the SegEM volume segmentation were extracted by collecting all voxels from the one-voxel boundary of the segmentation for which that pair of segments was present in the boundary’s 26-neighborhood. Then, all interface voxels for a given pair of segments were linked by connected components, and if multiple connected components were created, these were treated as separate interfaces. Interface components with a size of 150 voxels or less were discarded.

To define the subvolumes around an interface used for feature aggregation (Fig. 1k), we collected all voxels that were at a maximal distance of 40, 80 and 160 nm from any interface voxel and that were within either of the two adjacent segments of the interface. The interface itself was also considered as a subvolume yielding a total of 7 subvolumes for each interface.

### Feature calculation

11 3-dimensional image filters with one to 15 instances each (Table 1) were calculated as follows and aggregated over the 7 subvolumes of an interface using 9 summary statistics, yielding 3224 features per directed interface. Image filters were applied to cuboids of size 548x548x268 voxels, each, which overlapped by 72,72 and 24 voxels in x, y and z dimension, respectively, to ensure that all interface subvolumes were fully contained in the filter output.

Gaussian filters were defined by evaluating the unnormalized 3d Gaussian density function

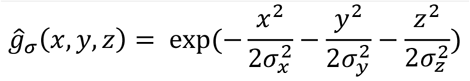

at integer coordinates 
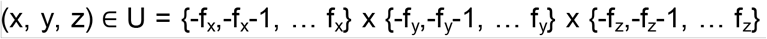
 for a given standard deviation σ = (σ_x_, σ_y_, σ_z_) and a filter size f = (f_x_, f_y_, f_z_) and normalizing the resulting filter by the sum over all its elements

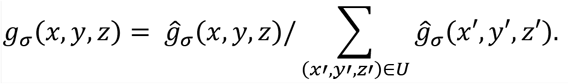

First and second order derivatives of Gaussian filters were defined as

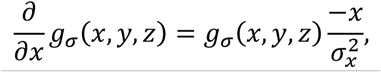

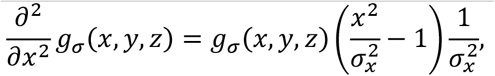

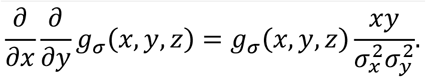

and analogously for the other partial derivatives. Normalization of g_σ_ and evaluation of derivatives of Gaussian filters was done on U as described above. Filters were applied to the raw data I via convolution (denoted by *) and we defined the image’s Gaussian derivatives as

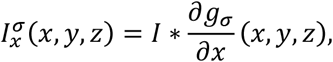

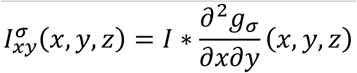

and analogously for the other partial derivatives.

Gaussian smoothing was defined as I*g_σ_.

Difference of Gaussians was defined as (I*g_σ_ - I*g_kσ_), where the standard deviation of the second Gaussian filter is multiplied element-wise by the scalar k.

Gaussian gradient magnitude was defined as

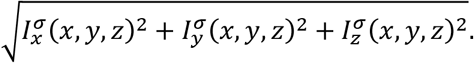

Laplacian of Gaussian was defined as

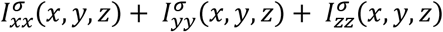

Structure tensor S was defined as a matrix of products of first order Gaussian derivatives, convolved with an additional Gaussian filter (window function) g_σw_:

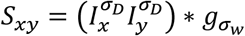

and analogously for the other dimensions, with standard deviation σ_D_ of the image’s Gauss derivatives. Since S is symmetric, only the diagonal and upper diagonal entries were determined, the eigenvalues were calculated and sorted by increasing absolute value.

The Hessian matrix was defined as the matrix of second order Gaussian derivatives:

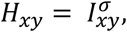

and analogously for the other dimensions. Eigenvalues were calculated as described for the Structure tensor.

The local entropy feature was defined as

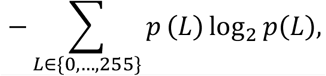

where p(L) is the relative frequency of the voxel intensity in the range {0, …, 255} in a given neighborhood U of the voxel of interest (calculated using the entropyfilt function in MATLAB).

Local standard deviation for a voxel at location (x, y, z) was defined by taking the maximum of 0 and

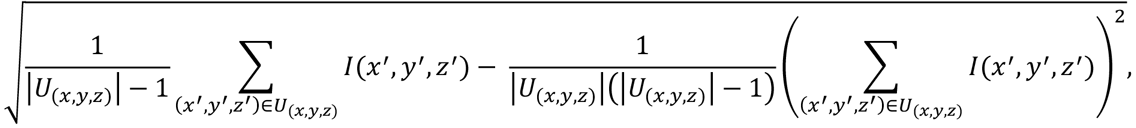

for the neighborhood U_(x,y,z)_ of location (x, y, z) with |U_(x,y,z)_| number of elements and calculated using MATLABs stdfilt function.

Sphere average was defined as the mean raw data intensity for a spherical neighborhood U_r_ with radius r around the voxel of interest, with

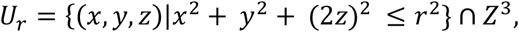

where Z^3^ is the 3 dimensional integer grid; x, y, z are voxel indices; z anisotropy was approximately corrected.

The intensity/variance feature for voxel location (x, y, z) was defined as

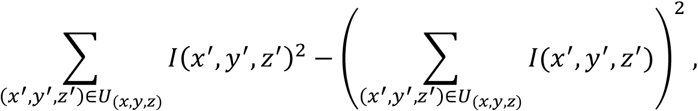

for the neighborhood U_(x,y,z)_ of location (x, y, z).

The set of parameters for which filters were calculated is summarized in Table 1. 11 shape features were calculated for the border subvolume and the two 160 nm-restricted subvolumes, respectively. For this, the center locations (midpoints) of all voxels of a subvolume were considered. Shape features were defined as follows: The number of voxel feature was defined as the total number of voxels in the subvolumes. The voxel based diameter was defined as the diameter of a sphere with the same volume as the number of voxels of the subvolumes. Principal axes lengths were defined as the three eigenvalues of the covariance matrix of the respective voxel locations. Principal axes product was defined as the scalar product of the first principal components of the voxel locations in the two 160 nm-restricted subvolumes. Voxel based convex hull was defined as the number of voxels within the convex hull of the respective subvolume voxels (calculated using the convhull function in MATLAB).

### Generation of training and validation labels

Interfaces were annotated by 3 trained undergraduate students using a custom-written GUI (in MATLAB, Suppl. Fig. 1a). A total of 40 non-overlapping rectangular volumes within the center 86 x 52 x 86 μm^3^ of the dataset were selected (39 sized 5.6 x 5.6 x 5.6 μm^3^ each and one of size 9.6 x 6.8 x 8.3 μm^3^). Then, all interfaces within these volumes were extracted as described above. Interfaces with a center of mass less than 1.124 µm from the volume border were not considered. For each interface, a raw data volume of size 1.6 x 1.6 x 0.7 μm^3^, centered on the center of mass of the interface voxel locations was presented to the annotator. When the center of mass was not part of the interface, the closest interface voxel was used. The raw data was rotated such that the second and third principal components of the interface voxel locations (restricted to a local surround of 15x15x7 voxels around the center of mass of the interface) defined the horizontal and vertical axes of the displayed images. First, the image plane located at the center of mass of the interface was shown. The two segmentation objects were transparently overlaid (Suppl. Fig. 1a) in separate colors (the annotator could switch the labels off for better visibility of raw data). The annotator had the option to play a video of the image stack or to manually browse through the images. The default video playback started at the first image. An additional video playback mode started at the center of mass of the interface, briefly transparently highlighted the segmentation objects of the interface, and then played the image stack in reverse order to the first plane and from there to the last plane. In most cases, this already yielded a decision. In addition, annotators had the option to switch between the 3 orthogonal reslices of the raw data at the interface location (Suppl. Fig. 1a). The annotators were asked to label the presented interfaces as non-synaptic or synaptic. For the synaptic label, they were asked to indicate the direction of the synapse (see Suppl. Fig. 1a). In addition to the annotation label interfaces could be marked as “undecided”. Interfaces were annotated by one annotator each. The interfaces marked as undecided were validated by an expert neuroscientist. In addition, all synapse annotations were validated by an expert neuroscientist, and a subset of non-synaptic interfaces was cross-checked. Together, 75,383 interfaces (1858 synaptic, 73,525 non-synaptic) were labeled this way. Of these, the interfaces from 8 label volumes (391 synaptic and 11906 non-synaptic interfaces) were used as validation set; the interfaces from the other 32 label volumes were used for training.

### SynEM classifier training and validation

The target labels for the binary, augmented and directed label sets were defined as described in the Results (Fig. 2c). We used boosted decision stumps (level-one decision trees) trained by the AdaBoostM1 (Freund & Schapire, 1997) or LogitBoost (Friedman et al., 2000) implementation from the MATLAB Statistical Toolbox (fitensemble). In both cases the learning rate was set to 0.1 and the total number of weak learners to 1500. Misclassification cost for the synaptic class was set to 100. Precision and recall values of classification results were reported with respect to the synaptic class. For validation, the binary label set was used, irrespective of the label set used in training. If the classifier was trained using the directed label set then the thresholded prediction for both orientations were combined by logical OR.

### Test set generation and evaluation

To obtain an independent test set disjunct from the data used for training and validation, we randomly selected a volume of size 512 × 512 × 256 voxels (5.75 × 5.75 × 7.17 μm^3^) from the dataset that contained no soma or dominatingly large dendrite. One volume was not used because of unusually severe local image alignment issues which are meanwhile solved for the entire dataset. The test volume had the bounding box [3713, 2817, 129, 4224, 3328, 384] in the dataset. First, the volume was searched for synapses (see Fig. 1d) in webKnossos (Boergens, Berning et al., under review) by an expert neuroscientist. Then, all axons in the volume were skeleton-traced using webKnossos. Along the axons, synapses were searched (strategy in Fig. 1e) by inspecting vesicle clouds for further potential synapses. Afterwards the expert searched for vesicle clouds not associated with any previously traced axon and applied the same procedure as above. In total, that expert found 335 potential synapses. A second expert neuroscientist used the tracings and synapse annotations from the first expert to search for further synapse locations. The second expert added 8 potential synapse locations. All 343 resulting potential synapses were collected and independently assessed by both experts as synaptic or not. The experts labeled 282 potential locations as synaptic, each. Of these, 261 were in agreement. The 42 disagreement locations (21 from each annotator) were re-examined jointly by both experts and validated by a third expert on a subset of all synapses. 18 of the 42 locations were confirmed as synaptic, of which one was just outside the bounding box. Thus, in total, 278 synapses were identified. The precision and recall of the two experts in their independent assessment with respect to this final set of synapses was 93.6%, 94.6% (expert 1) and 97.9%, 98.9% (expert 2), respectively.

Afterwards all shaft synapses were labeled by the first expert and proofread by the second. Subsequently, the synaptic interfaces were voxel-labeled to be compatible with the method by Becker et al. This initial test set comprised 278 synapses, of which 36 were labeled as shaft/inhibitory.

Next, all interfaces between pairs of segmentation objects in the test volume were extracted as described above. Then, the synapse labels were assigned to those interfaces whose border voxels had any overlap with one of the 278 voxel-labeled synaptic interfaces. Afterwards, these interface labels were again proof-read by an expert neuroscientist. Finally, interfaces closer than 160 nm from the boundary of the test volume were excluded to ensure that interfaces were fully contained in the test volume. The final test set comprised 235 synapses out of which 31 were labeled as shaft/inhibitory. With this we obtained a high-quality test set providing both voxel-labeled synapses and synapse labels for interfaces, to allow the comparison of different detection methods.

For the calculation of precision and recall, a synapse was considered detected if at least one interface that had overlap with the synapse was detected by the classifier (TPs); a synapse was considered missed if no overlapping interface of a given synapse was detected (FNs); and a detection was considered false positive (FP) if the corresponding interface did not overlap with any labeled synapse.

### Comparison to previous work

The approach of Becker et al., 2012 was evaluated using the implementation provided in Ilastik (Sommer et al., 2011). This approach requires voxel labels of synapses. We therefore first created training labels: an expert neuroscientist created sparse voxel labels at interfaces between pre-and postsynaptic processes and twice as many labels for non-synaptic voxels for five cubes of size 3.4 x 3.4 x 3.4 μm^3^ that were centered in five of the volumes used for training SynEM. Synaptic labels were made for 115 synapses (note that the training set in Becker et al., 2012 only contained 7-20 synapses). Non-synaptic labels were made for two training cubes first. The non-synaptic labels of the remaining cubes were made in an iterative fashion by first training the classifier on the already created synaptic and non-synaptic voxel labels and then adding annotations specifically for misclassified locations using Ilastik. Eventually, non-synaptic labels in the first two training cubes were extended using the same procedure.

For voxel classification all features proposed in (Becker et al., 2012) and 200 weak learners were used. The classification was done on a tiling of the test set into cubes of size 256x256x256 voxels (2.9 x 2.9 x 7.2 μm^3^) with a border of 280 nm around each tile. After classification, the borders were discarded, and tiles were stitched together. The classifier output was thresholded and morphologically closed with a cubic structuring element of three voxels edge length. Then, connected components of the thresholded classifier output with a size of at least 50 voxels were identified. Synapse detection precision and recall rates were determined as follows: A ground truth synapse (from the final test set) was considered detected (TP) if it had at least a single voxel overlap with a predicted component. A ground truth synapse was counted as a false negative detection if it did not overlap with any predicted component (FN). To determine false positive classifications, we evaluated the center of the test volume (shrunk by 160 nm from each side to 484 x 484 x 246 voxels) and counted each predicted component that did not overlap with any of the ground truth synapses as false positive detection (FP). For this last step, we used all ground truth synapses from the initial test set, in favor of the Becker et al. classifier.

For comparison with (Kreshuk et al., 2014) the same voxel training data as for (Becker et al., 2012) was used. The features provided by Ilastik up to a standard deviation of 5 voxels for the voxel classification step were used. For segmentation of the voxel probability output map the graph cut segmentation algorithm of Ilastik was used with label smoothing ([1, 1, 0.5] voxel standard deviation), a voxel probability threshold of 0.5 and graph cut constant of λ = 0.25. Objects were annotated in five additional cubes of size 3.4 x 3.4 x 3.4 μm^3^ that were centered in five of the interface training set cubes different from the one used for voxel prediction resulting in 299 labels (101 synaptic, 198 non-synaptic). All object features provided by Ilastik were used for object classification. The evaluation on the test set was done as for (Becker et al., 2012).

### Pairwise connectivity model

The neuron-to-neuron connection recall was calculated assuming an empirical distribution p(n) of the number of synapses n between connected neurons given by published studies (see Supp. Table 2, Feldmeyer et al., 1999, Feldmeyer et al., 2002, Feldmeyer et al., 2006, Frick et al., 2008, Markram et al., 1997). We further assumed the number of retrieved synapses given by a binomial model with retrieval probability given by the synapse classifier recall R_s_ on the test set:

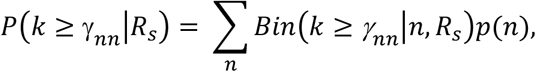

Where γ_nn_ is the threshold on the number of synapses between a neuron pair to consider it as connected (see Fig. 3a). This equates to the neuron-to-neuron recall: R_nn_ = P(k ≥ γ_nn_ | R_s_).

To compute the neuron-to-neuron precision, we first calculated the expected number of false positive synapse detections (FP_s_) made by a classifier with precision P_s_ and recall R_s_:

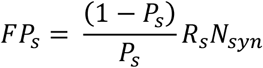

where N_syn_ is the total number of synapses in a dataset calculated from the average number of synapses per connected neuron pair <n_syn_> times the number of connected neuron pairs N_con_ and c_r_ is the connectivity ratio given by N_con_/N^2^ with N the number of neurons in the connectome.

We then assumed that these false positive synapse detections occur randomly and therefore are assigned to one out of N^2^ possible neuron-to-neuron connections with a frequency FP_s_/N^2^.

We then used a Poisson distribution to estimate the number of cases in which at least γ_nn_ FP_s_ synapses would occur in a previously zero entry of the connectome, yielding a false positive neuron-to-neuron connection (FP_nn_).

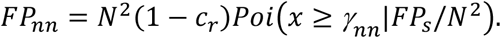

Finally, the true positive detections of neuron-to-neuron connections in the connectome TP_nn_ are given in terms of the neuron-to-neuron connection recall R_nn_ by

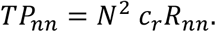

Together, the neuron-to-neuron connection precision P_nn_ is given by

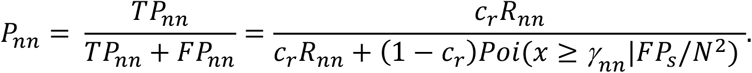

The connectivity ratio was set to c_r_ = 0.2 (Feldmeyer et al., 1999).

### Local connectome

For determining the local connectome (Fig. 4) between 100 pre-and 100 postsynaptic processes, we used 100 axonal skeleton tracings (traced at 1 to 5-fold redundancy) and 100 dendrite skeleton tracings. All volume objects which overlapped with any of the skeleton nodes were detected and concatenated to a given neurite volume. Then, all interfaces between pre-and postsynaptic processes were classified by SynEM. The area of each interface was calculated as in (Berning et al., 2015) and the total area of all contacts between all neurite pairs was calculated (Fig. 4c). To obtain the binary connectome C_bin_ (Fig. 4d), we applied the SynEM score threshold θ_nn_ (Fig. 2e, Table 3), and the connectome threshold γ_nn_ = 1. To obtain a weighted connectome C_w_ (Fig. 4e), we re-analyzed all interfaces between neurites determined to be connected in the binary connectome. This time, the SynEM score threshold θ_s_ was applied (see Fig. 2e, Table 3). Detected synaptic interfaces were clustered using hierarchical clustering (single linkage, distance cutoff 1,500 nm) if the interfaces were between the same pre-and postsynaptic objects. The connection probability as a function of total contact area between neurites (Fig. 4 g) was calculated as the fraction of neurite-to-neurite connections with at least A_c_ total contact area that was synaptically connected.

## AUTHOR CONTRIBUTIONS

Conceived and initiated the project: MH; supervised the project: MH and PvdS; Developed algorithms, implemented algorithms, analyzed data: BS; provided segmentations and contributed to algorithm development: MB; provided EM data: KMB; provided expert synapse annotations: AG; wrote the paper: MH and BS.

## ACKNOWLEDGEMENTS

We thank Jan Gleixner for first test experiments on synapse detection and fruitful discussions in an early phase of the project, Alessandro Motta for comments on the manuscript, and Raphael Jacobi, Raphael Kneissl, Athanasia Natalia Marahori, and Anna Satzger for data annotation.

## REFERENCES

Bartol, T. M., Bromer, C., Kinney, J., Chirillo, M. A., Bourne, J. N., Harris, K. M., & Sejnowski, T. J. (2015). Nanoconnectomic upper bound on the variability of synaptic plasticity. Elife, 4, e10778. doi:10.7554/eLife.10778

Becker, C., Ali, K.,Knott, G., & Fua, P. (2012). Learning context cues for synapse segmentation in EM volumes. Paper presented at the International Conference on Medical Image Computing and Computer-Assisted Intervention.

Becker, C., Ali, K., Knott, G., & Fua, P. (2013). Learning context cues for synapse segmentation. IEEE Trans Med Imaging, 32(10), 1864-1877. doi:10.1109/TMI.2013.2267747

Berning, M., Boergens, K. M., & Helmstaedter, M. (2015). SegEM: Efficient Image Analysis for High-Resolution Connectomics. Neuron, 87(6), 1193-1206. doi:10.1016/j.neuron.2015.09.003

Bock, D. D., Lee, W. C., Kerlin, A. M., Andermann, M. L., Hood, G., Wetzel, A. W., Reid, R. C. (2011). Network anatomy and in vivo physiology of visual cortical neurons. Nature, 471(7337), 177-182. doi:10.1038/nature09802

Braitenberg, V., & Schüz, A. (1998). Cortex: statistics and geometry of neuronal connectivity: Springer Science & Business Media.

Briggman, K. L., Helmstaedter, M., & Denk, W. (2011). Wiring specificity in the direction-selectivity circuit of the retina. Nature, 471(7337), 183-188. doi:10.1038/nature09818

Colonnier, M. (1968). Synaptic patterns on different cell types in the different laminae of the cat visual cortex. An electron microscope study. Brain Res, 9(2), 268-287.

Denk, W., & Horstmann, H. (2004). Serial block-face scanning electron microscopy to reconstruct three-dimensional tissue nanostructure. PLoS Biol, 2(11), e329. doi:10.1371/journal.pbio.0020329

Eberle, A. L., Mikula, S., Schalek, R., Lichtman, J., Knothe Tate, M. L., & Zeidler, D. (2015). High-resolution, high-throughput imaging with a multibeam scanning electron microscope. J Microsc, 259(2), 114-120. doi:10.1111/jmi.12224

Feldmeyer, D., Egger, V., Lubke, J., & Sakmann, B. (1999). Reliable synaptic connections between pairs of excitatory layer 4 neurones within a single 'barrel' of developing rat somatosensory cortex. J Physiol, 521 Pt 1, 169-190.

Feldmeyer, D., Lubke, J., & Sakmann, B. (2006). Efficacy and connectivity of intracolumnar pairs of layer 2/3 pyramidal cells in the barrel cortex of juvenile rats. J Physiol, 575(Pt 2), 583-602. doi:10.1113/jphysiol.2006.105106

Feldmeyer, D., Lubke, J., Silver, R. A., & Sakmann, B. (2002). Synaptic connections between layer 4 spiny neurone-layer 2/3 pyramidal cell pairs in juvenile rat barrel cortex: physiology and anatomy of interlaminar signalling within a cortical column. J Physiol, 538(Pt 3), 803-822.

Freund, Y., & Schapire, R. E. (1997). A decision-theoretic generalization of on-line learning and an application to boosting. Journal of Computer and System Sciences, 55(1), 119-139. doi:DOI 10.1006/jcss.1997.1504

Frick, A., Feldmeyer, D., Helmstaedter, M., & Sakmann, B. (2008). Monosynaptic connections between pairs of L5A pyramidal neurons in columns of juvenile rat somatosensory cortex. Cereb Cortex, 18(2), 397-406. doi:10.1093/cercor/bhm074

Friedman, J., Hastie, T., & Tibshirani, R. (2000). Additive logistic regression: A statistical view of boosting. Annals of Statistics, 28(2), 337-374. doi:DOI 10.1214/aos/1016218223

Gray, E. G. (1959). Axo-somatic and axo-dendritic synapses of the cerebral cortex: an electron microscope study. J Anat, 93, 420-433.

Hayworth, K., Kasthuri, N., Schalek, R., & Lichtman, J. (2006). Automating the collection of ultrathin serial sections for large volume TEM reconstructions. Microscopy and Microanalysis, 12(S02), 86-87.

Helmstaedter, M. (2013). Cellular-resolution connectomics: challenges of dense neural circuit reconstruction. Nat Methods, 10(6), 501-507. doi:10.1038/nmeth.2476

Helmstaedter, M., Briggman, K. L., & Denk, W. (2011). High-accuracy neurite reconstruction for high-throughput neuroanatomy. Nat Neurosci, 14(8), 1081-1088. doi:10.1038/nn.2868

Helmstaedter, M., Briggman, K. L., Turaga, S. C., Jain, V., Seung, H. S., & Denk, W. (2013). Connectomic reconstruction of the inner plexiform layer in the mouse retina. Nature, 500(7461), 168-174. doi:10.1038/nature12346

Hua, Y., Laserstein, P., & Helmstaedter, M. (2015). Large-volume en-bloc staining for electron microscopy-based connectomics. Nat Commun, 6, 7923. doi:10.1038/ncomms8923

Huang, G. B., Scheffer, L. K., & Plaza, S. M. (2016). Fully-Automatic Synapse Prediction and Validation on a Large Data Set. arXiv preprint arXiv:1604.03075.

Kasthuri, N., Hayworth, K. J., Berger, D. R., Schalek, R. L., Conchello, J. A., Knowles-Barley, S., Lichtman, J.W. (2015). Saturated Reconstruction of a Volume of Neocortex. Cell, 162(3), 648-661. doi:10.1016/j.cell.2015.06.054

Knott, G., Marchman, H., Wall, D., & Lich, B. (2008). Serial section scanning electron microscopy of adult brain tissue using focused ion beam milling. J Neurosci, 28(12), 2959-2964. doi:10.1523/JNEUROSCI.3189-07.2008

Kreshuk, A., Funke, J., Cardona, A., & Hamprecht, F. A. (2015). Who is talking to whom: synaptic partner detection in anisotropic volumes of insect brain. Paper presented at the International Conference on Medical Image Computing and Computer-Assisted Intervention.

Kreshuk, A., Koethe, U., Pax, E., Bock, D. D., & Hamprecht, F. A. (2014). Automated detection of synapses in serial section transmission electron microscopy image stacks. PLoS One, 9(2), e87351. doi:10.1371/journal.pone.0087351

Kreshuk, A., Straehle, C. N., Sommer, C., Koethe, U., Cantoni, M., Knott, G., & Hamprecht, F. A. (2011). Automated detection and segmentation of synaptic contacts in nearly isotropic serial electron microscopy images. PLoS One, 6(10), e24899. doi:10.1371/journal.pone.0024899

Lee, W. C., Bonin, V., Reed, M., Graham, B. J., Hood, G., Glattfelder, K., & Reid, R. C. (2016). Anatomy and function of an excitatory network in the visual cortex. Nature, 532(7599), 370-374. doi:10.1038/nature17192

Markram, H., Lubke, J., Frotscher, M., Roth, A., & Sakmann, B. (1997). Physiology and anatomy of synaptic connections between thick tufted pyramidal neurones in the developing rat neocortex. J Physiol, 500(Pt 2), 409-440.

Merchan-Perez, A., Rodriguez, J. R., Gonzalez, S., Robles, V., Defelipe, J., Larranaga, P., & Bielza, C. (2014). Three-dimensional spatial distribution of synapses in the neocortex: a dual-beam electron microscopy study. Cereb Cortex, 24(6), 1579-1588. doi:10.1093/cercor/bht018

Mikula, S., Binding, J., & Denk, W. (2012). Staining and embedding the whole mouse brain for electron microscopy. Nat Methods, 9(12), 1198-1201. doi:10.1038/nmeth.2213

Mikula, S., & Denk, W. (2015). High-resolution whole-brain staining for electron microscopic circuit reconstruction. Nat Methods, 12(6), 541-546. doi:10.1038/nmeth.3361

Mishchenko, Y., Hu, T., Spacek, J., Mendenhall, J., Harris, K. M., & Chklovskii, D. B. (2010). Ultrastructural analysis of hippocampal neuropil from the connectomics perspective. Neuron, 67(6), 1009-1020. doi:10.1016/j.neuron.2010.08.014

Pallotto, M., Watkins, P. V., Fubara, B., Singer, J. H., & Briggman, K. L. (2015). Extracellular space preservation aids the connectomic analysis of neural circuits. Elife, 4. doi:10.7554/eLife.08206

Sommer, C., Straehle, C., Kothe, U., & Hamprecht, F. A. (2011). Ilastik: Interactive Learning and Segmentation Toolkit. 2011 8th Ieee International Symposium on Biomedical Imaging: From Nano to Macro, 230-233.

White, E. L., & Peters, A. (1993). Cortical modules in the posteromedial barrel subfield (Sml) of the mouse. J Comp Neurol, 334(1), 86-96. doi:10.1002/cne.903340107

